# Robustness of cortical distortion metrics to unmodelled OPM errors

**DOI:** 10.64898/2026.11.17.767363

**Authors:** Alberto Mariola, Stephanie Mellor, James J. Bonaiuto, Tim M. Tierney, John Ashburner, Yaël Balbastre, José David López, Gareth R. Barnes

## Abstract

The estimate of cortical current flow from magnetic fields measured outside the head is an ill-posed problem. Recent work using head casts to minimise co-registration errors in conventional cryogenic magnetoencephalography (MEG) has shown that ground truth anatomy can be used to judge between different inversion schemes and metrics. Inversion algorithms or metrics that are insensitive to anatomical distortion are unlikely to reflect correct assumptions about the underlying current distribution.

Here we aim to identify constraints when using this approach to validate current flow estimated from recordings made with optically pumped MEG (OP-MEG). To do so, we used simulations contaminated by different sources of (typically unmodelled) error. For example: error in the estimate of the true sensor positions or orientations and the effects of unmodelled gain changes.

In accordance with previous work, with no unmodelled errors, distortion was minimized when the simulation and the inversion assumptions matched. We then quantified the amount of error (in mm) relative to the true cortical mesh that one might expect in empirical recordings (given gain, position and orientation errors). We found that the approach was tolerant to independent sensor position errors of less than 2 mm (per axis); orientation errors of less than 5 degrees; fixed array displacements of less than 10 degrees and to random sensor gain errors of less than 10%. Additionally, we tested this approach using two empirical OPM datasets. We found that source reconstructions based on both minimum norm and beamformer assumptions were sensitive to the underlying anatomy.

## Introduction

Optically Pumped Magnetometers (OPMs) represent a cryogen-free alternative to superconducting quantum interference device (SQUID) sensors for the measurement of neural activity. OPMs can be flexibly placed in wearable structures (or scanner-casts), and have been used in personalised (Boto et al., 2017) and generic (Budker & Romalis, 2007; Boto et al., 2017; Tierney et al., 2019; Hill et al., 2020; Brookes et al., 2022) helmets to record magnetoencephalography (MEG).

The emergence of lightweight, wearable OPM arrays allows researchers to move away from classical static experimental settings towards more naturalistic paradigms (Seymour et al., 2021; Mellor et al., 2023; O’Neill et al., 2025; Mellor et al., 2026). This has many advantages but makes the comparison with cryogenic MEG or supine fMRI-compatible experimental designs challenging. This results in a disconnect between imaging modalities and re-opens the search for objective ways to quantify the validity of MEG source reconstruction pathways.

Source reconstruction is an ill-posed inversion problem (i.e., magnetic fields measured from outside the head could derive from an infinite combination of neural sources) which requires the introduction of constraints based on biologically and physically plausible assumptions. While different source reconstruction schemes are based on different assumptions (Mosher et al., 2003), they each provide subjectively plausible estimates of neuronal current flow.

In the present work, we leverage realistic anatomical distortion algorithms (Balbastre et al., 2018; Ashburner et al., 2019) to introduce a framework that aims to objectively evaluate any MEG or EEG analysis pathway. The main assumption behind this framework is that, since the external magnetic fields measured by M/EEG are necessarily linked to anatomy (i.e., intracellular primary current flow is bound to pyramidal neurons), then the use of distorted anatomical models (e.g. distorting cortical meshes obtained from participants’ MRI) should lead to suboptimal source estimates. This validation through anatomy becomes less ill-posed the more data are collected as the anatomy remains constant across all recorded data for the same subject (Barnes et al., 2006; Stevenson et al., 2014; López et al., 2017; Little et al., 2018; López et al., 2026). Most recently, López et al. (2026) showed that surrogate cortical meshes, which approximate true cortical anatomy, can be used to quantify the performance of source reconstruction algorithms and metrics of fit (using cryogenic MEG data). These surrogate anatomies are generated by means of diffeomorphic shape modelling (Balbastre et al., 2018; Ashburner et al., 2019), an algorithm that parametrizes individual brain anatomy with a compact set (∼100) of Gaussian latent variables. In the present work, we aim to test the robustness of the method to typical sources of unmodelled errors in OP-MEG data.

We use simulations to test the effect of unmodelled errors during OPM recordings on this framework. To do this, we first simulated OPM data without errors and then attempted to reconstruct these data under deviations from the true OPM recording set-up (i.e., the data remains the same but the forward model changes). First, we randomly shifted sensor positions and attempted to reconstruct these data. Second, we manipulated the orientation of the three axes (all by the same amount, in the same direction) associated with each OPM sensor (triaxial). Third, we randomly rotated all sensors (and axes) by the same amount around the head to simulate rigid (fixed helmet) rotation errors. Fourth, we assessed the influence of random gainerrors applied directly to the lead field matrices. Finally, we made use of empirical OPM data to test our approach using real task recordings.

In summary, we find that this method is sensitive to unmodelled geometrical errors. Additionally, we show that the source reconstructions from our two OP-MEG empirical datasets were sensitive to the anatomical distortion; but the method could not distinguish between different inversion schemes.

## Methods

We wanted to test the robustness of the anatomical distortion framework that was previously employed with cryogenic MEG and head-casts (López et al., 2026) to the typical unmodelled errors one might encounter using OPMs. To do this, we used the method outlined in López and colleagues (2026) but added typical errors specific to current OPM systems.

The following outlines how we first distort the anatomy to create surrogate cortical surfaces which deviate, by varying degree (quantified in millimetres), from the truth. This creates a parameter space over which we can test inversion algorithms and metrics. The logic being that an algorithm (or metric) which fits equally well to the distorted as to the true anatomy is suboptimal. Following López et al. (2026), we use negative free energy as the fit metric (Friston et al., 2007).

We then simulate a single source on the true cortical surface and reconstruct it using appropriate and inappropriate algorithms (as we know the source covariance structure, we know which algorithms should and should not work).

The data always remained the same, but we selectively distorted the lead fields linking the cortex to the sensors to simulate unmodelled errors due to imperfect OPM sensor arrays.

For zero unmodelled error added, we expect the approach to identify the correct imaging assumptions (or algorithm); the aim is to see how large these unmodelled errors can be before the approach stops working.

Finally, we used the standard approach to see if we could distinguish between two commonly used algorithms for reconstructing empirical MEG data derived from OPM sensors. Here, modelling errors are intrinsic to the experimental setup and unknown, rather than explicitly simulated.

### Diffeomorphic Modelling

The specific diffeomorphic modelling algorithms that we employed in this study are the ones used by López et al. (2026), based on the work of Balbastre and colleagues (2018). The main idea behind diffeomorphic shape modelling is that any individual anatomical structure can be represented by applying a smooth, invertible (diffeomorphic) transformation to a shared anatomical template (Miller et al., 1993).

In this modelling paradigm, the transformations are defined by initial velocity fields, and differences between shapes are measured based on how smooth these fields are. Once the average template has been established, a Bayesian residual component analysis (Bishop, 1998; Kalaitzis & Lawrence, 2012) is used to analyse the variability in the velocity fields. This method, a variation of principal component analysis, accounts for and removes known sources of structured variation (in this case, smoothness), focusing instead on meaningful shape differences.

This means that each anatomical structure can be characterized using a compact set of latent variables that capture its most significant variations (Zhang & Fletcher, 2015; Balbastre et al., 2018; Ashburner et al., 2019). The model enables the generation of novel anatomical samples by drawing random latent variables from a standard Gaussian distribution.

### Parameter Space

To account for the normal brain shape variation, the model uses 100 eigenvectors. This means that the shape of each individual brain (and corresponding cortical mesh) can be defined by a 100-element vector (**z**). To generate a series of distorted versions of the original cortical mesh, we manipulated this individual vector (**z**). Following López and colleagues (2026), since the first seven components/elements mostly represent information associated with size and volume of the brain, we only distorted the remaining 93 components (8-100). Each distortion trajectory is defined as a 93-dimensional vector with values of +1 or −1. This means that each component will move either from {−*Z* to +*Z*} or {+*Z* to −*Z*}, wherein *Z* encodes the normal distortion range. A discrete index *j* codes for distortion from the original mesh. The movement of *j* from 1 to *N* (i.e., number of points/discrete meshes on the distortion trajectory) traverses a mesh from maximum deviation in one direction to maximum deviation in the other; passing close to the true mesh at the middle of the range. Specifically, here we used *Z* = 3 and *N* = 17 (the 18th mesh corresponding to the original one). Since each individual distortion trajectory is defined by a random seed (which defines the 93-element distortion trajectory), in this work we used eight different seeds for the simulation and 64 for the empirical data (in general the more seeds the better). Full mathematical details about the generative shape model and of the specific instantiation of cortical mesh distortions can be found in both Balbastre and colleagues (2018) and López and colleagues (2026).

### Simulated Data

OP-MEG data were simulated using SPM25 (Tierney et al., 2025) by means of the *spm_opm_sim* function. Specifically, we simulated one dataset containing 56 OPM triaxial sensors (168 channels of data) and 100 trials lasting 500 ms (with −150 ms of pre-stimulus activity) sampled at 375 Hz.

Sensor positions were based on the personalised scanner-cast of the participant reported in the Empirical Data section of this manuscript. Similarly, cortical mesh, inner skull, outer skull and skin over which data were inverted were those of the same participant. For each of the 100 trials, we then simulated a single 20 Hz sinusoidal dipolar source between 0 and 350 ms, a dipolar moment of 10 nAm and a sensor-level signal-to-noise ratio of −20 dB at the average location of the human’s left motor cortex (MNI Coordinates: *x* = −40, *y* = −18, *z* = 48).

#### Sensor Errors

To assess how specific sensor errors might affect the performance of our cortical distortion framework, we kept the underlying source and anatomy constant but artificially introduced a series of errors to the forward model of our simulated system across a series of dimensions: sensor position errors, orientation errors, rigid rotation errors and gain errors.

The application of unmodelled errors to the forward models followed Mellor and colleagues (2024). Four types of error were considered:

1. <u>Position errors</u>. These errors can be due to mischaracterization of the location of the active sensor position on the head (for example OPMs attached flexibly and independently to the scalp). These errors were applied by independently shifting each sensor by a specific distance extracted from a zero-mean Gaussian distribution with a given standard deviation along each axis (x, y or z). Importantly, errors were consistent across all three channels within each sensor, but different across sensors. Standard deviation values of 0, 0.5, 1, 2, 5 and 10 mm per axis were used. These within axis deviations correspond to absolute sensor displacements of 0, 0.86, 1.73, 3.46, 8.66, and 17.32 mm respectively.
2. <u>Rigid rotation errors</u>. These errors might be due to a rigid array of sensors (housed in a helmet for example) which are imprecisely registered to the scalp. We initially selected a random rotation axis (x, y or z) around which we rotated the entire helmet (akin to MEG co-registration errors in which participants’ heads move with respect to a fixed helmet). Practically, this was done by building a rotation matrix that was applied to all sensors. Rotation angles were extracted from a zero-mean Gaussian distribution with standard deviations of 0, 1, 2, 5, 10 and 20°.
3. <u>Orientation errors</u>. These might be due to flexible arrays or sub-optimally characterised sensing axes. To characterize these errors, we moved the three axes (x, y, z channels) of each sensor by means of a rotation matrix encoding a shift in their orientation (keeping the axes orthogonal). Once again, all three axes of the same sensors were rotated consistently, but differently across sensors. Rotation angles were extracted from a zeromean Gaussian distribution with standard deviations of 0, 5, 10 and 20°,
4. <u>Gain errors</u>. These can arise due to non-zero fields across the OPM sensors (Iivanainen et al., 2019). Gain errors were created by multiplying the lead field matrices of the simulated OPM data with a vector of gain coefficients. Gain coefficients were extracted from a unit-mean Gaussian distribution with standard deviations 0, 5, 10, 15, 20% and they were randomly adjusted over sensors.

All errors were applied independently (i.e., we did not apply two or more error categories to our simulated OPM data at the same time). For each error level we drew eight possible values from the distribution (each error level defines a standard deviation over channels); and for each of these simulations we ran eight different seeds of distortion.

For example, we chose eight position errors based on a standard deviation of 2 mm; for each of these position error values we reconstructed onto 18 different cortical surfaces to find the one that gave rise to maximum negative free energy (Friston et al., 2007). This gave a single metric of distortion in millimetres. This was repeated eight times, each time using a different seed (or a different set of 18 surfaces). We did this for two inversion schemes: Empirical Bayes Beamformer (EBB - Belardinelli et al., 2012) and Minimum Norm (Hämäläinen & Ilmoniemi, 1994). This means that there were at least 8×8×18×2 (=2304) inversions per level of unmodelled error.

### Empirical Data

#### Acquisition

To test the framework empirically, we first applied it to OP-MEG data recorded from a single healthy participant (male, 57-years old, right-handed) taking part in two tasks. The first task consisted of a series of 102 continuous and self-paced button presses that the participant was asked to produce in intervals approximately 3 seconds apart. The second task, from Mellor and colleagues (2026), was a roving mismatch negativity paradigm (auditory oddball) based on Garrido and colleagues (2008). Full details regarding the task can be found in both Seymour et al. (2021) and Mellor et al. (2023, 2026). In summary, the participant was exposed to a series of 70 ms long tones (5 ms rise and fall times) at different randomly selected frequencies between 500 Hz and 800 Hz in steps of 50 Hz. In total, the task comprised of 571 tones (trials), with 80 deviant tones. The inter-stimulus interval was 500 ms (without stimulus jitter). Stimuli were presented via PsychoPy (Peirce, 2009), through MEG-compatible ear tubes. Crucially, for the main analyses, we ignored the nature of the tone (standard vs deviant) and we focused on source reconstruction (and related cortical mesh distortion) of auditory evoked responses to all tones. Our participant completed both tasks across two different conditions: either while being seated or while walking around the centre of the OPM Magnetic Shielded Room (MSR; Magnetic Shields Ltd, UK) located in the Department of Imaging Neuroscience at University College London. For the present paper, we only focus on the seated condition. The room has internal dimensions of 438 cm × 338 cm × 218 cm and comprises two inner layers of 1 mm mu-metal, a 6 mm copper layer, and then two external layers of 1.5 mm mu-metal.

OP-MEG data for both tasks were recorded by means of a Neuro-1 system (QuSpin, Louisville, CO) composed of 56 triaxial sensors (168 channels) and they were sampled at 1500 Hz. Both datasets were recorded in closed-loop mode. OPM sensors positions were co-registered with participants’ anatomy by means of a participant-specific, 3D-printed scanner-cast (Chalk Studios Ltd., UK) that was designed and built from the participant’s structural MRI scan (Meyer et al., 2017 - see MRI Acquisition).

#### Preprocessing

Data were preprocessed in MATLAB R2022a using SPM25 (Tierney et al., 2025) and a series of custom scripts (see Data Availability section). Initially, the power spectral density of OP-MEG data was visually examined to detect and manually remove bad channels. For the auditory task, we removed 46 channels (out of 168), for the button press task, we removed 50 channels. Data were then high-pass filtered at 2 Hz, band-stop filtered (48-52 Hz) and low-pass filtered at 70 Hz. All filters were 5th order Butterworth filters applied bidirectionally to achieve zerophase shift. To remove environmental magnetic interference, we used adaptive multipole models (AMM; Tierney et al., 2024) with default spatial parameters (internal harmonics order: 9; external harmonic order: 2; correlation limit: 1). After temporal and spatial filtering, data were downsampled to 375 Hz. For the auditory task, continuous data were then epoched into 450 ms segments centred around the onset of each tone (-100 ms to 350 ms) whereas button press data were segmented into 2400 ms epochs (-1200 ms to 1200 ms) to avoid overlap between consecutive trials (source reconstruction was then computed on the evoked button press response between −200 and 300 ms). In total, the auditory task contained 571 trials, whereas the participant completed 102 button presses. All trials were visually inspected, and bad trials were removed by means of the Fieldtrip’s *spm_eeg_ft_artefact_visual* function (Oostenveld et al., 2011) implemented in SPM. In total, we removed 52 epochs for the auditory oddball paradigm and 11 trials for the button press task. Source reconstruction and corresponding anatomical distortions were then based on the average of all trials of both tasks respectively.

### MRI Acquisition

Full details of MRI data acquisition can be found in Bonaiuto et al. (2018). In summary, the protocol featured a quantitative multi-parameter mapping (MPM) approach, which included three multi-echo 3D FLASH scans with different weightings. These scans were RF- and gradient-spoiled and provided whole-brain coverage at an isotropic resolution of 800 µm. The protocol also included additional calibration measurements to correct for RF transmit field inhomogeneities (Lutti et al., 2010; Lutti et al., 2012; Callaghan et al., 2015). A 32-channel head coil was employed during this protocol to enhance signal-to-noise ratio.

### Source Reconstruction and Model Comparison

The MRI of our single participant was used to generate the cortical mesh and forward model needed to invert both empirical and simulated OPM data. Specifically, we obtained the individual cortical surface by segmenting the T1-weighted MRI using FreeSurfer’s *recon-all* function (Fischl, 2012). We used a white matter cortical surface model consisting of 33,596 vertices (current dipoles), with orientation defined by the local surface normal vectors (constituting our source model). We used the Single Shell (Nolte, 2003) forward model.

Lead fields (i.e., gain matrices) were recomputed once for each mesh (*n* = 18) and error level, error seeds and distortion trajectory seeds (*n* = 128; four-or more-error levels, eight error seeds, and eight and 64 distortion seeds for simulated and empirical data respectively) resulting in a total of more than 2,304 gain matrices.

Following Phillips et al. (2002) and López and colleagues (2014), both simulated and empirical data were reduced to 16 temporal modes (via singular value decomposition). Additionally, a spatial projector was also applied to the empirical data over the lead field matrices of both datasets (99 components for both auditory oddball and button press tasks). Crucially, within each dataset and distortion seed, the same spatial modes were used before inversion on all cortical meshes.

To explain both our simulated and empirical OP-MEG data as current flow constrained to each surface on the distortion trajectory, we employed two source reconstruction algorithms defined by different modelling assumptions (encoded by their source covariance):

- The Minimum Norm algorithm (referred to here as Identically Independently Distributed Noise; IID - Hämäläinen & Ilmoniemi, 1994), which assumes all the potential sources to be both i) equiprobable and ii) independent/uncorrelated from each other (i.e., the source covariance is equal to a scaled identity matrix).
- The Empirical Bayes Beamformer (EBB) (Belardinelli et al., 2012), a Bayesian formulation of the Linearly Constrained Minimum Variance (LCMV) beamformer (Drongelen et al., 1996; Van Veen et al., 1997), in which the (diagonal) source covariance is estimated empirically from the data via spatial filtering and forms the prior. This prior is then weighted with the noise (assumed to be a scaled identity matrix).

For both empirical and simulated data, we used negative variational free energy (Friston et al., 2007; also known as the evidence lower bound - ELBO) as the optimization and model fit metric (Stevenson et al., 2014; López et al., 2017; Little et al., 2018; López et al., 2026). Free energy is a strictly positive quantity, in which zero indicates a perfect model fit, hence we use negative free energy so that the optimal model returns a maximal value.

### Statistical Significance

To assess whether the mean distances (from the true cortical mesh) associated with the maximum negative free energy value obtained across the different simulation settings and empirical datasets were statistically indistinguishable from chance, we built non-parametric null distributions for both categories. The null distribution of distances (i.e., distortion estimates) was obtained by applying the anatomical distortion framework to source inversions that incorporated null lead field matrices in which rows (encoding specific sensors) were randomly shuffled. In this way, the to-be-inverted data remained unchanged but any biophysical link to the true anatomy (determined by the lead fields) was broken (López et al., 2026).

We repeated this procedure on the same 64 distortion seeds employed for the original empirical data (eight seeds for simulations) and across eight lead field shuffling randomisation seeds thereby generating 512 distances per inversion scheme (64 distances for simulations). We then constructed a null distribution of mean distances per inversion scheme (EBB and IID) by averaging 64 randomly selected distance values (eight for simulations) 1,000 times. We then assessed whether the observed average distance from the original brain (i.e., distortion) was lower than the 95th percentile of the combination of both distributions. Finally, a Welch’s t-test was used to test the distortion values differences between inversion schemes across simulated data with no-error and empirical datasets.

## Results

The main goal of this study was to assess the fundamental constraints that our anatomical distortion framework might face when reconstructing the current flow associated with both simulated and empirical OP-MEG data. We assessed how moving through the space of potential surrogate cortical meshes affected the source reconstruction estimates of simulated OPM data influenced by different types and levels of unmodelled errors. Then, we applied our anatomical distortion framework to real OPM data recorded from one participant undergoing an auditory oddball task and a button press paradigm.

### Simulated Data

A single sinusoidal source (20 Hz) was placed in the left motor cortex of the cortical mesh. We first attempted to localise this source via two inversion schemes: EBB and IID, reflecting matching (spatially focal source activity) and ill-matching (sources assumed to be independent, equally likely, and identically distributed) assumptions, respectively. The inversion procedure was then repeated for additional 17 distorted cortical meshes (with a mean vertex-to-vertex distance range from the original mesh of: 0.5 – 4 mm) across eight possible distortion seeds (different distortion trajectories). Figure 1A shows that for EBB (with assumptions matching the simulation), the negative free energy cost function generally peaked at the true cortical mesh (or zero distortion) for seven out of eight possible distortion trajectories. Conversely, for IID (illmatching assumptions) the cost-function did not follow the underlying anatomical distortion gradient. The mean distortion across the eight peaks (in the cost function) was lower than 95% of the null distribution (Figure 1B and 1C - red dotted line) for EBB (*M* = 0.17, *SE* = 0.17, *p* < .001) but not for IID (*M* = 2.46, *SE* = 0.37, *p* = .072), with the EBB mean distortion value being significantly lower than the IID one (*t*(9.83) = −5.58, *p* < .001). Note the similarity between the 95% threshold for IID and EBB permutations. For clarity, we use the joint distribution to calculate a single 95% threshold in subsequent simulation figures.

**Figure 1.**
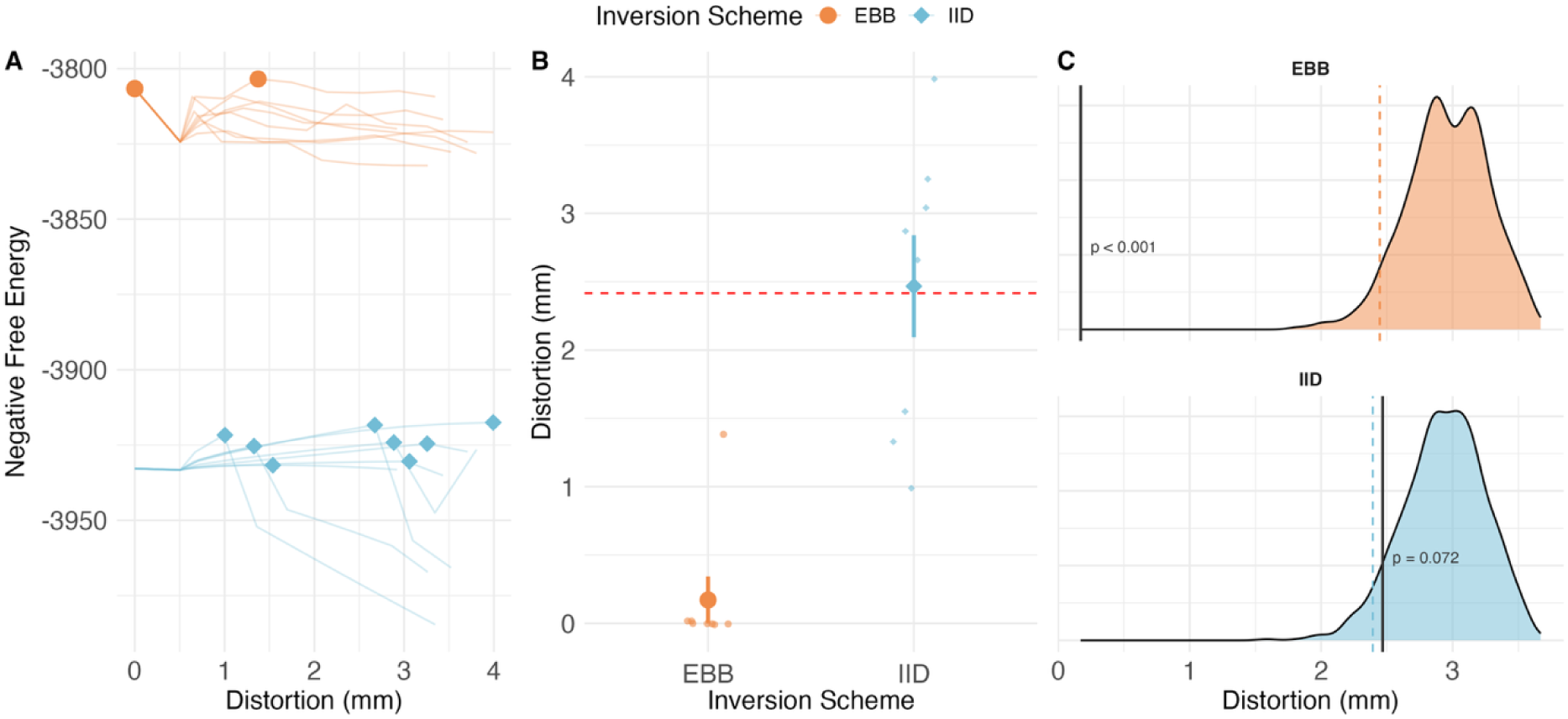
– **Simulated Data. A)** Line plot showing the negative free energy associated with reconstructions onto the different cortices as distortion is increased for eight different trajectories across two different algorithms (EBB, orange and IID, blue). Circles and diamonds mark the peak metric (signifying most likely underlying cortical mesh) for EBB and IID respectively. For EBB, across seven out of eight possible distortion trajectories, the most likely cortical mesh corresponded to the original one (overlapping circles). **B**) Scatterplot showing the distance (in mm) from the true cortical mesh associated with the maximum free energy value for both EBB (orange circle) and IID (blue diamond). Each of the smaller points represents the computed distance value associated with each distortion seed. Large, bold points correspond with the average distance from the true cortical mesh for the two source inversion schemes. Error bars show the standard error of the mean. **C)** Inversion scheme specific null distributions of distortion values (distances from true cortical mesh) obtained by permuting the lead fields. Each distribution is composed of 1000 permuted mean distortion values (EBB in orange; IID in blue). Orange (EBB) and blue (IID) dashed lines represents the lowest 95th percentile values of each distribution. The red dashed line represents the lowest 95th percentile (Figure 1B) combining the two distributions. Black solid lines represent the observed mean distortion value (with corresponding p-value).

#### Simulation Errors

We then explored how different error types and levels would influence the reconstruction performance of both inversion schemes and how this interacted with the cortical manifold distortion approach (Figure 2). Briefly, we found the distortion framework to be sensitive to position errors of 2 mm per axis, orientation errors of 5 degrees, rigid rotations of 10 degrees, and lead field gain errors of 10%.

**Figure 2.**
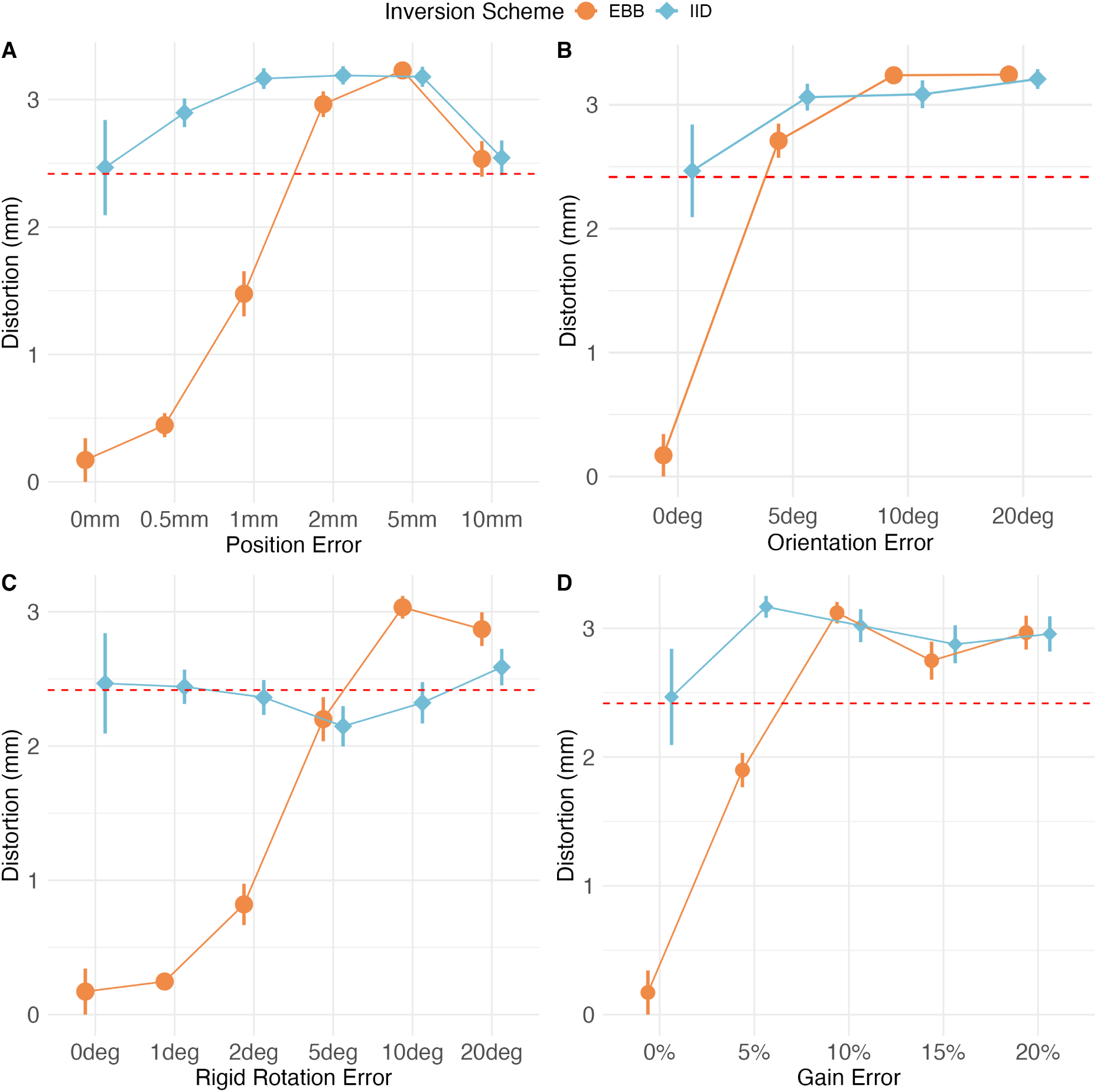
– **Simulated Data - Errors.** Line plots showing the distance (in mm) from the true cortical mesh associated with the maximum negative free energy value for both EBB (orange circles) and IID (blue diamonds) across different error types (one per panel) and levels. Bold points correspond with the average distance from the true cortical mesh for the two source inversion schemes. Error bars show standard error of the mean, whereas the red dashed line corresponds to the lowest 95th percentile obtained from the combination of the permuted null distributions of the simulated data (without errors). **A**. Distortion estimates as a function of position error. These position error values are set over 3 axes of space (x, y, and z), so with absolute displacements of 0, 0.87, 1.73, 3.46, 8.66, and 17.32 mm respectively (values multiplied by √3 to account for the three dimensions). **B**. Distortion estimates as a function of orientation error (identical across three sensor axes, with channels remaining orthogonal). **C**. Distortion estimates as a function of rigid rotation error (e.g., where the entire sensor array rotated as a rigid body). **D**. Distortion estimates as a function of random gain errors.

### Empirical Data

We then applied the same approach to real OP-MEG data collected from a single participant completing two experimental tasks: an auditory oddball paradigm (Figure 3) and a button press task (Figure 4). Figures 3 and 4 show subjectively plausible current source estimates for both schemes. For the auditory task (Figure 5A-B-C) there was significant dependence of the free energy cost-function on the true anatomy for both EBB (*M* = 2.52, *SE* = 0.19, *p* < .001) and IID (*M* = 2.36, *SE* = 0.19, *p* < .001). Similarly, for the button press task (Figure 5D-E-F), the mean distance associated with the highest free energy values was significantly lower than the permuted ones for both EBB (*M* = 3.18, *SE* = 0.12, *p* = 0.03) and IID (*M* = 3.03, *SE* = 0.12, *p* = 0.002). However, in contrast with the pattern observed during simulations, there was no significant difference between the average distortion values observed for EBB and IID across both the auditory oddball task (*t*(125.28) = 0.53, *p* = 0.59) and spontaneous button presses one (*t*(125.58) = 0.80, *p* = 0.42).

**Figure 3.**
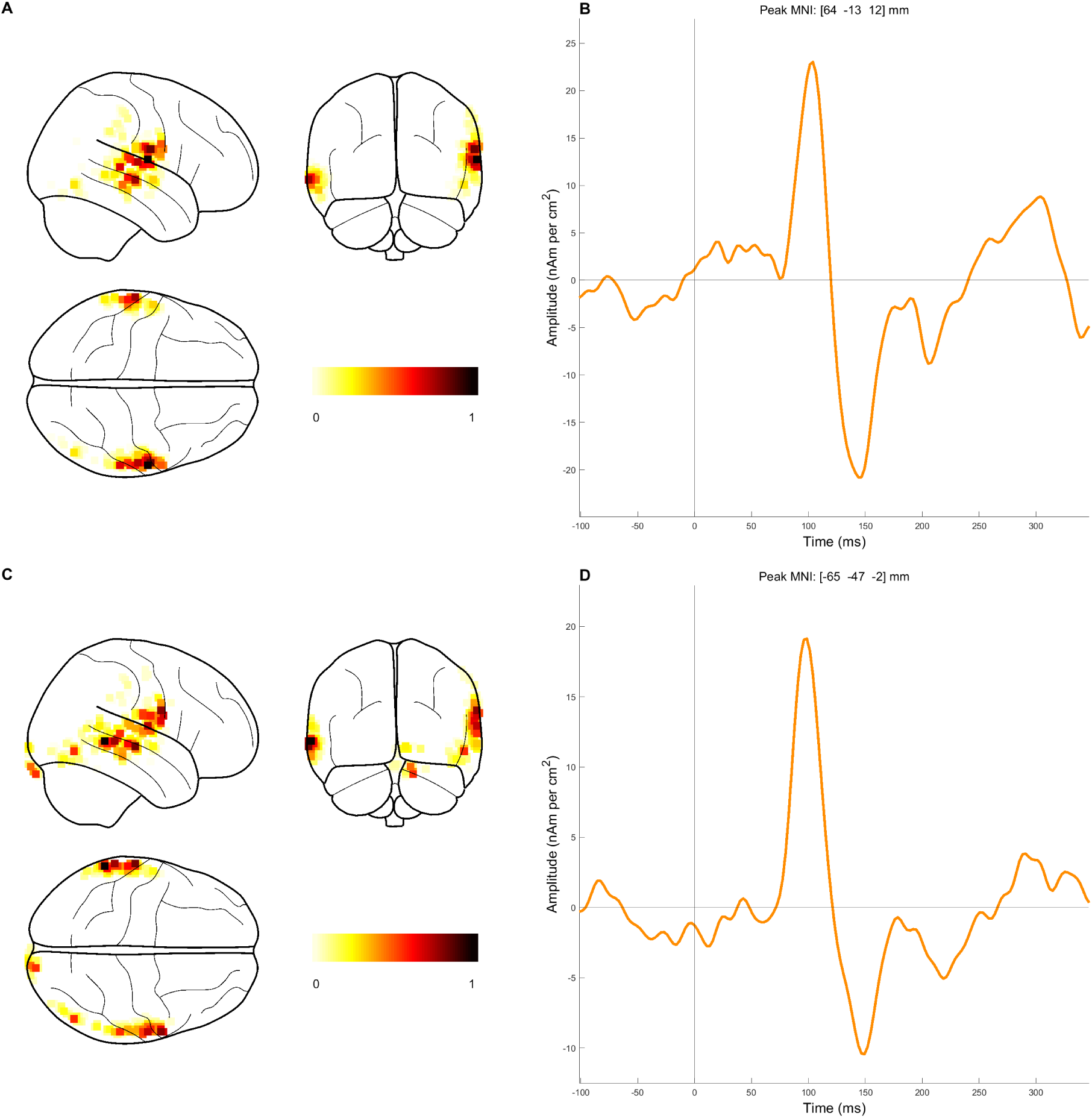
– **Auditory Evoked Response - Source Reconstruction.** Source reconstruction of the average auditory evoked response obtained from the auditory oddball paradigm via EBB (Panel A and B) and IID (Panel C and D). Panel A and C contain two glass brain views showing voxels associated with the 99th percentile of normalised power over the entire period of interest. Colour bars indicate normalised values (0-1) associated with increasing power between the 99th and 100th percentile. Panels B and D show the time course of the reconstructed response corresponding to the peak vertex (indicated in MNI coordinates).

**Figure 4.**
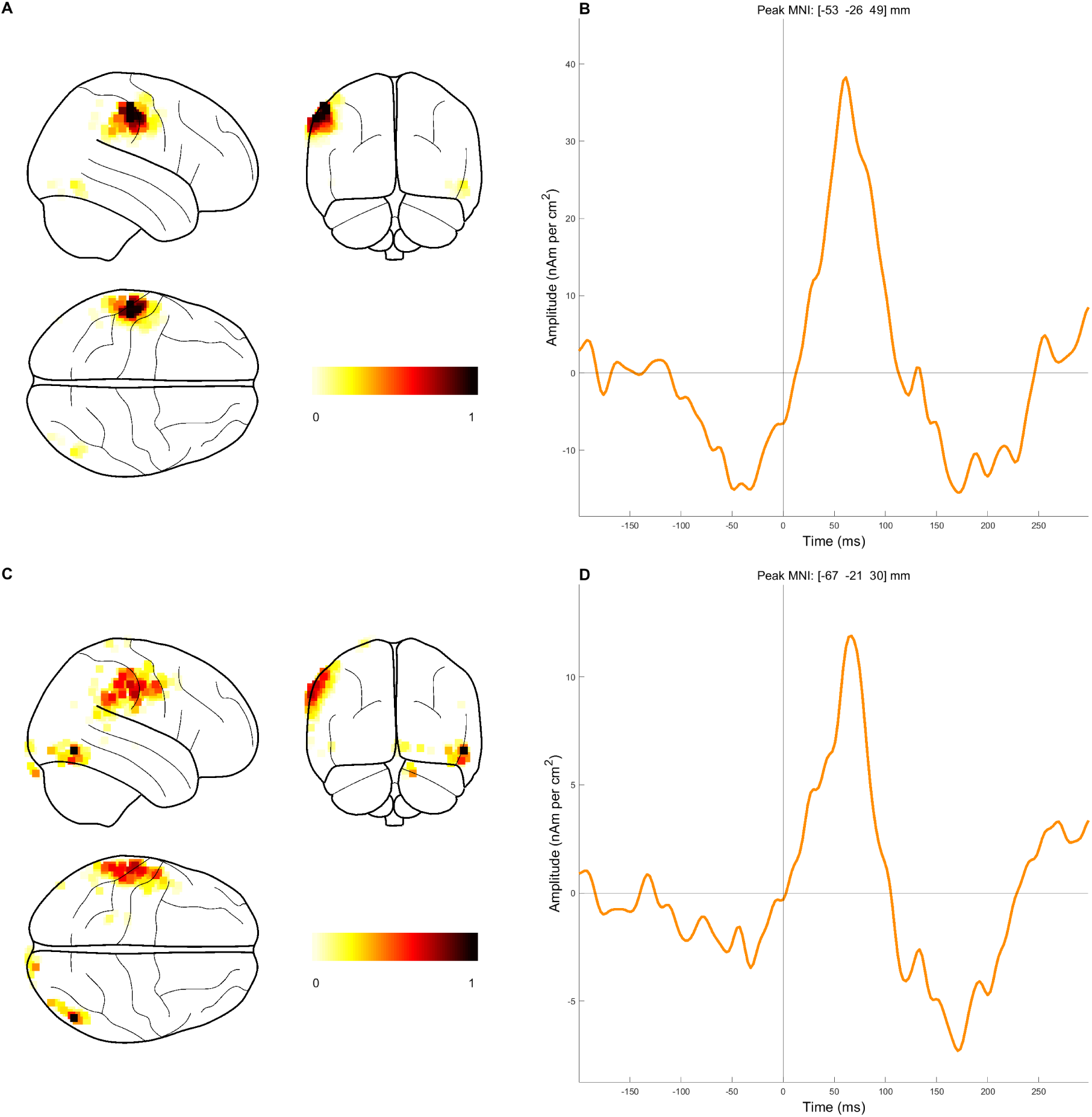
– **Motor Evoked Response - Source Reconstruction.** Source reconstruction of the average motor evoked response obtained from the auditory oddball paradigm via EBB (Panel A and B) and IID (Panel C and D). Panel A and C contain two glass brain views showing voxels associated with the 99th percentile of normalised power over the entire period of interest. Colour bars indicate normalised values (0-1) associated with increasing power between the 99th and 100th percentile. Panels B and D show the time course of the reconstructed response corresponding to the peak vertex (indicated in MNI coordinates).

**Figure 5.**
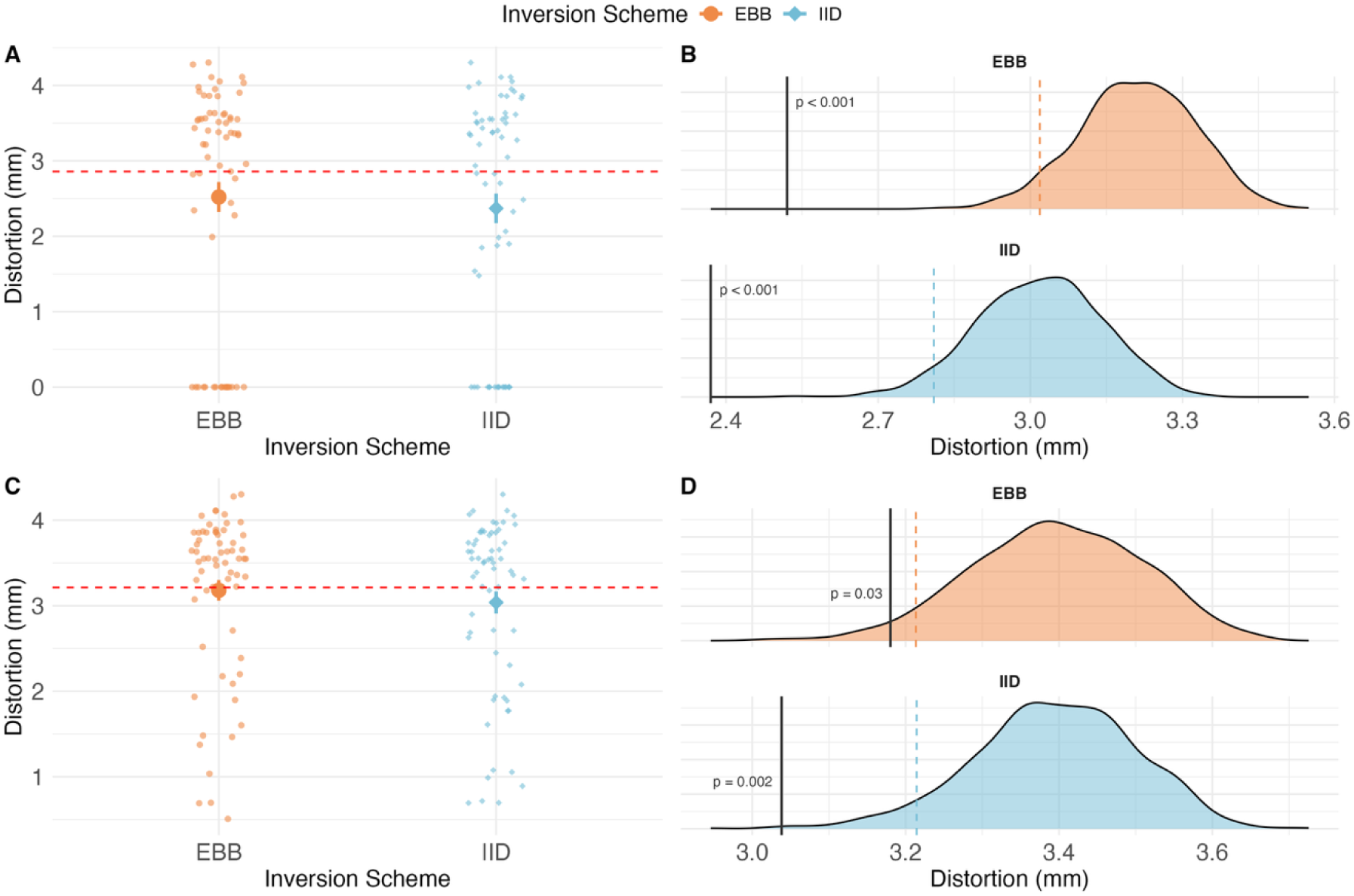
– **Empirical Data. A)** Auditory Evoked Response - Scatterplot showing the distance (in mm) from the true cortical mesh associated with the maximum negative free energy value for both EBB (orange) and IID (blue) as compared to the null data (dotted). Each of the small points represents the computed distance value associated with each distortion seed. Bold points correspond to the average distance from the true cortical mesh for the two source inversion schemes. Vertical line represents the standard error of the mean. **B)** Auditory Evoked Response - Inversion scheme specific null distributions of distortion values (distances from true cortical mesh) obtained by permuting the lead fields. Each distribution is composed of 1,000 permuted mean distortion values (EBB in orange; IID in blue). Orange (EBB) and blue (IID) dashed lines represents the lowest 95th percentile values of each distribution. The red dashed line represents the lowest 95th percentile (Figure 5A) combining the two distributions. Black solid lines represent the observed mean distortion value (with corresponding p-value). Panel **C** and **D** represent the corresponding results for the motor evoked response.

## Discussion

In this paper we expanded on recent work of López et al. (2026), in which anatomy defines ground truth, from cryogenic to optically pumped MEG. We examined the sensitivity of this approach to unmodelled errors specific to typical OPM set-ups as well as its performance on empirical OP-MEG data.

### Simulated Data

In the ideal case of no errors, we replicated the findings of López and colleagues (2026). The approach was sensitive to the true anatomy only in the case where inversions assumptions matched the simulated data.

However, by introducing unmodelled simulation errors, we were able to explore the tolerance of the approach. We found that for this framework to work, random gain errors had to be less than 10%, individual channel orientation had to be known within 5 degrees; channel position within 2 mm per axis (or 3.46 mm in any direction) and rigid rotation errors (due to rigid helmets for example) to less than 10 degrees.

We have shown that this method is sensitive to the complete OPM pathway: from recording to reconstruction assumptions. Veridical sensors’ characterization of the set-up is required to make a distinction between reconstruction algorithms, metric or forward models. However, once that has been established (e.g. which algorithm, forward model to use), this requirement (as the data can now be trusted) can be relaxed. That said, the requirement for precise knowledge of OPM geometry for source reconstruction has been noted by others (Zetter et al., 2018; Hill et al., 2020). For example, Hill et al. (2020) found that co-registration errors of more than 2 degrees/ mm per axis for both rigid helmets (corresponding to our rigid rotation case) and flexible caps (corresponding to our position error case) caused source localisation errors of around 2 mm (with the error growing to ranges of 10 mm to 16 mm when the co-registration errors reached 10 mm). In this context, recent developments in sensor localisation technologies that can precisely recover sensors’ positions and axes (channels) orientations such as the HALO system (QuSpin, Louisville, CO) or tailored external coils (Iivanainen et al., 2022; Hill et al., 2025) represent a fundamental advancement in the field. Some commercial systems now also offer built-in autolocalisation, in which the sensors localise themselves using either coils embedded in the helmet (HEDscan OPM system - FieldLine Medical, Louisville, CO; Alem et al., 2023) or by acting as magnetic field emitters and receivers (MAG4Health, Grenoble, France).

### Empirical Data

The functional estimates obtained empirically appear plausible across both experimental cases (Figures 3 and 4) and all the inversion schemes assessed by means of our anatomical distortion approach significantly outperformed the results obtained by the null case (Figure 5). This indicates that this new generation of sensors, coupled with traditional reconstruction algorithms, shows a stronger dependence on cortical anatomy than one would expect by chance (Figure 5). Importantly, these findings are largely consistent with the ones obtained by López and colleagues (2026) with cryogenic MEG.

The distortion framework could not, however, disambiguate between the two inversion schemes (EBB and IID) in either task (Figure 5). In contrast, in both cases, there were significant differences between the observed negative free energy estimates (Figure S1 – Supplementary Materials). That is, from a mathematical modelling perspective, EBB (with greater negative free energy in both cases) is the most likely model to describe brain function. The distortion metric goes further to demand anatomic specificity but did not suggest that these current estimates were any more likely than those from IID. López et al. (2026) reported a similar pattern of results. Empirically, we minimised geometric errors using an individualized scanner-cast. At the time of recording, we were only able to precisely locate the sensor casing (i.e., we did not have an external calibration device) and so sensor position and orientation errors due to individual sensor engineering were not quantified. These errors have been shown to be on the order of ∼2mm in position and, on average, 10 degrees per axis in orientation from that expected ideally (Hill et al., 2025). This meant that the empirical measurement axes were likely not only misaligned but also not orthogonal to one another. In other words, the geometric errors due to the divergence between the sensor casing and sensor itself could account for the reduction in performance in real data. Likewise, the gain errors estimated by Hill and colleagues (2025) were of the order of 1-2% per axis.

We should stress that here no knowledge of (cortical) functional, but precise sensor and anatomical (e.g., position error < 2 mm, rigid rotation error < 10 degrees, individual orientation error < 5 degrees etc.), prior knowledge is required. In the cryogenic case, the precise anatomical knowledge was possible due to head cast recordings (in which the subject’s head is completely constrained). In future, OPMs fitted to an individualized head cast can be worn comfortably with well-defined geometry relative to the brain for long periods of time.

### Limitations

While our results highlight the robustness and flexibility of the anatomical distortion approach when applied to both simulated and empirical OP-MEG data, there are several methodological considerations.

First, each seed determines a specific distortion trajectory; it may be, however, that this anatomical distortion occurs in a region that is not functionally active. There are two possible mitigating approaches. The first is to summarize changes across multiple seeds (assuming at least one of them will distort the region of active cortex) as done here using 64 different seeds or distortion trajectories. The second (used by Little et al., 2018) is to use more (and objectively identified) data through unsupervised portioning of the whole dataset into self-similar states through, for example, Hidden Markov Modelling (this means that ideally more functional areas will be included in the analysis). Future research might combine latest developments in the state-space modelling field (Vidaurre et al., 2018; Gohil et al., 2022; Vidaurre et al., 2025) with realistic distortion of participants’ cortical manifolds. This would pave the way for fully end- to-end data-driven source reconstruction pipelines that would also allow for the analysis of complex resting-state and naturalistic datasets.

The anatomical precision required necessitates the acquisition of a recent MRI scan for each participant. At present, this necessarily limits the use of this approach to studies which include the collection of a T1-weighted image and that ideally features the production of a personalised scanner-cast for each participant. This is clearly expensive and time-consuming, with the subsequent analysis rather intensive. One issue with the current analysis could be that the cortical surface used was made based on the subject’s anatomy collected eight years ago whilst supine; while OP-MEG data was acquired during recent upright recording conditions. Because of this, we cannot exclude that both the temporal and postural discrepancy of the two datasets could have influenced the quality of our functional estimates and subsequent performance of our distortion framework. Likewise, segmentation errors may constrain the accuracy to which the true surface, and its diffeomorphic surrogates, can be represented (Seiger et al., 2018; Beelen et al., 2020; Larson & Oguz, 2022).

Furthermore, it is important to highlight that for the present work we used the white matter surface (in accordance with Stevenson et al., 2014 and López et al., 2026), as we expect the large pyramidal neurons to be the dominant source of current flow (Sherman et al., 2016). However, our framework could be easily extended to the use of the pial surface or any intermediate cortical surface resulting from the interpolation of white matter and pial surfaces (e.g., Szul et al., 2026). This may be a useful research avenue given recent investigations around layer and frequency specific cortical dynamics associated with different sensory and cognitive process across both cryogenic (Troebinger et al., 2014; Bonaiuto et al., 2018; Szul et al., 2026) and OP-MEG simulations (Helbling, 2025).

We adopted a Single Shell (Nolte, 2003) forward model for all our inversions (consistent with López et al., 2026). This model is computationally efficient and has been shown to outperform single shell Boundary Element Method (BEM) approaches (Stenroos et al., 2014). Future work could investigate the effect that more complex Finite Element Method (FEM) that precisely characterise different compartments (e.g., skull, CSF) could have on the solution.

Additionally, it is worth noting that, following Mellor and colleagues (2024), we applied each unmodelled error independently in our simulations. This choice was motivated by the need to precisely characterise each source of error separately, as well as by the high computational cost of simulating all potential interactions between error types and levels. However, some sources of error are likely to interact during empirical recordings. For example, as noted by Mellor et al. (2024), if an environmental magnetic field gradient affected the entire sensor array, sensor gain and position would become highly correlated. Future research should examine the impact of these interactions on cortical distortion metrics.

Relatedly, cross-axis projection error (CAPE) (Borna et al., 2022) might have reduced the performance of our approach on empirical OP-MEG data. CAPE arises when a remnant static field, particularly along the sensor’s laser propagation axis, is combined with signal fields on multiple axes. This rotates each sensor’s sensitive axis, alters its gain, and introduces frequency-dependent phase errors. Because the remnant field differs between sensors and changes as sensors move through field gradients (for example, in wearable systems), these errors are sensor-specific and potentially time-varying, unlike the independent, static errors we simulated. Future studies could extend the suite of simulated errors to include CAPE, for example by coupling sensor orientation and gain errors to a modelled remnant field.

The main point of this paper was to identify the tolerances on sensor placement and precision. We were able to show that the method works empirically on a single-subject case study collected during multiple tasks. This may motivate future studies to make use the anatomical distortion framework across other OP-MEG datasets recorded in different experimental settings.

## Conclusion

We examined how an objective metric of distance from ground truth could be applied to OP-MEG recordings. For OPMs one might anticipate not only tasks than cannot be validated by an alternative supine scanner (like fMRI); but also, much longer recording times (more data per subject) and increasingly complex naturalistic experimental designs. We found that it should be possible within certain sensor tolerances. The objective is to create a global validation of any non-invasive functional estimate based on known anatomy.

## Ethics

The empirical data reported in the present paper were part of the sample and experimental paradigm reported in Mellor and colleagues (2026). The study protocol was in full accordance with the Declaration of Helsinki, and all participants gave written informed consent after being fully informed about the purpose of the study. The study protocol, participant information, and form of consent, were approved by the UCL Research Ethics Committee (reference number 5833/001).

## Data and Code Availability

The simulated and anonymised empirical OPM datasets will be made available via Zenodo upon publication. The anatomical distortion codebase is part of the development version of SPM (https://github.com/spm/spm). The full analysis pipeline can be found on GitHub (https://github.com/langestroop/opm-mesh-dist).

## Author Contributions

AM, GRB, JDL, YB - conception. AM, SM, TMT, GRB, JDL, JJB, YB, JA- coding. All authorsmanuscript writing and editing.

## Funding

JDL was funded by the Colombian MinCiencias project contract 1222-852-69927.

YB was funded in part by the MRC and Spinal Research Charity through the ERA-NET Neuron joint call (MR/R000050/1), and in part by a fellowship from the Royal Society (NIF∖R1∖232460). This research was supported by the Discovery Research Platform for Naturalistic Neuroimaging funded by the Wellcome (226793/Z/22/Z) and MRC equipment grant (MR/X012409/1). We would like to thank the UCL OP-MEG group for their helpful and constructive comments. TT was funded by an Epilepsy Research Institute fellowship (FY2101).

## Declaration of Competing Interest

The authors declare no competing interests.

## Declaration of the use of AI

We did not use AI in writing this paper. However, Claude Sonnet 5.5 (Anthropic) was used to help developing the code needed to generate Figure 3 as well as the structuring and organization of the project’s codebase on the GitHub repository.

## Notes

https://github.com/langestroop/opm-mesh-dist

### Competing Interest Statement

The authors have declared no competing interest.

