## Supplementary Materials for "Robustness of cortical distortion metrics to unmodelled OPM errors"

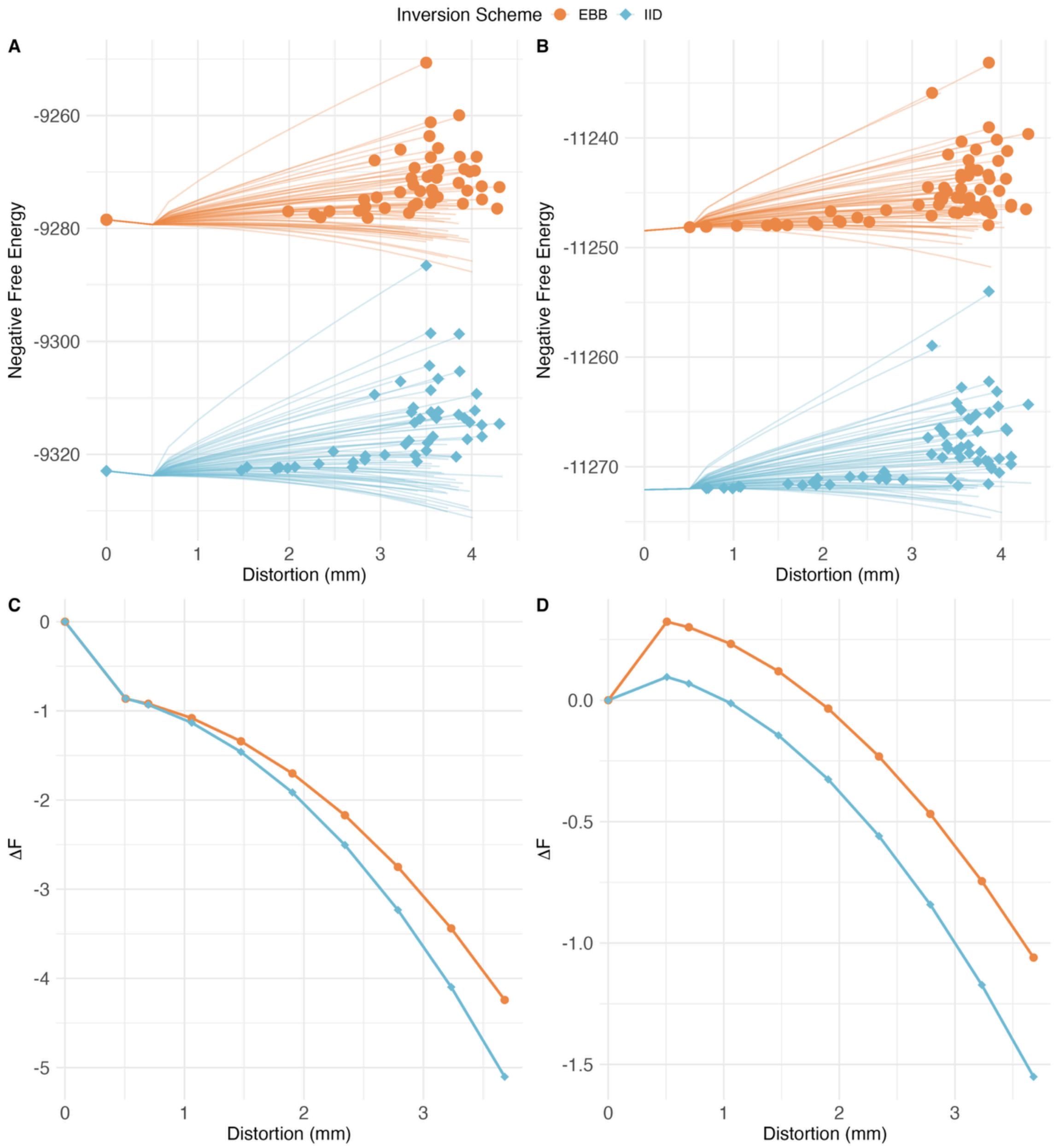


**Figure S1 – Empirical Data. A)** Auditory Evoked Response - Line plot showing the negative free energy associated with reconstructions onto the different cortices as distortion is increased for 64 different trajectories across two different algorithms (EBB, orange and IID, blue). Circles and diamonds mark the peak metric (signifying most likely underlying cortical mesh) for EBB and IID respectively. Note that symbols are overlaid and distortion 0 has 17 points for both EBB and IID. Panel **B** represents the corresponding result for the motor evoked response. **C)** Auditory Evoked Response - Line plot showing the average delta negative free energy (ΔF) as a function of distance from the true cortical mesh for both EBB (orange) and IID (blue). Average delta negative free energy represents the mean difference in negative free energy computed between its corresponding value for the original mesh and each discrete step (bold points) of the distortion trajectory across all distortion seeds. Panel **D** represents the corresponding result for the motor evoked response.
